# Diffusion MRI of the Unfolded Hippocampus

**DOI:** 10.1101/2020.04.07.029850

**Authors:** Uzair Hussain, Jordan DeKraker, Nagalingam Rajakumar, Corey A. Baron, Ali R. Khan

## Abstract

The hippocampus is implicated in numerous neurological disorders and the ability to detect subtle or focal hippocampal abnormalities earlier in disease progression could significantly improve the treatment of patients. Ex vivo studies with ultra-high field have revealed that diffusion MRI (dMRI) can reveal microstructural variations within the hippocampal subfields and lamina, and may also be sensitive to intra-hippocampal pathways. However, translation to lower resolution in vivo dMRI studies of the hippocampus is challenging due to its complicated geometry. One novel way to overcome some of these obstacles is by transforming the usual Cartesian coordinates in an MRI image to coordinates that are crafted to curve themselves according to the complicated geometry of the hippocampus. This procedure allows us to virtually unfold the hippocampus into a thin sheet. In this work, we introduce an algorithm to map diffusion MRI data to this sheet, allowing us to overcome the difficulties associated with the hippocampus’ complicated geometry. We demonstrate how our method can be readily integrated into existing implementations of traditional tractography methods and how it leads to enhancements in the resulting tracts. Further, our results on high quality in vivo dMRI acquisitions show that unfolding the hippocampus leads to a more anatomically plausible modelling of the connectivity of the hippocampus as probed by probabilistic tractography, revealing key elements of the polysynaptic pathway and anterior-posterior connectivity gradients.

## 1 Introduction

The hippocampus is a heavily studied structure of the brain. It plays a central role in learning processes, memory, and spatial navigation. It is also involved in major disease states. In Alzheimer’s disease (AD) the hippocampus is one of the first regions of the brain to undergo damage which significantly affects memory and cognition [1]. Hippocampal sclerosis (HS) is a significant lesion found in patients with temporal lobe epilepsy [2], [3]. HS presents itself with different patterns of cell loss and clinicopathologic studies hint towards a correlation of HS subtypes with post-surgical outcomes [4]. Improved identification of HS subtypes could allow for precise clinical decision-making leading to better post-surgical outcomes.

Although the hippocampus is not identical to the neocortex, it is similar and is frequently classified as archicortex given that it is a phylogenetically ancient cortical structure [5]. It has three-layers making its laminar structure discernible from the neighbouring multi-layered entorhinal cortex. The hippocampus can be partitioned into three distinct sub-regions, the dentate gyrus (DG), the hippocampus proper, and the subiculum (Sub). The hippocampus proper consists of three subfields, called CA1, CA2 and CA3. In cross sections, the CA subfields and the DG form two interlocked “C” shapes. The different areas of the hippocampus show specialization for different functions, for pattern separation and completion a key role is played by the DG and CA3, respectively, and CA2 is more involved in social memory [6], [7]. According to a standard connectional view [8], the main inputs to the hippocampus come from the entorhinal cortex, which is the source of the perforant pathway (PP) that projects to all the sub-regions of the hippocampus. The PP is so-called because it perforates the Sub. Within the hippocampus, the polysynaptic pathway is a collection of fibers that connects the sub-regions of the hippocampus in a sequential and unidirectional manner. Briefly stated, it starts in the DG where mossy fiber pathways aim CA3, from CA3 the Schaffer collaterals project to CA1 which finally projects to the Sub. Output from the hippocampus emerges from CA1 and the Sub, which then flows towards the entorhinal cortex.

Macroscopically, the hippocampus has a complex curved geometry and it bears some resemblance to the horns of a ram. This complex geometry and its small size makes it a challenge for most in-vivo imaging techniques. Nevertheless, dMRI based parcellation in combination with functional MRI has uncovered the existence of distinct functional networks within the hippocampus [9]. Deterministic tractography has been used to create detailed maps of the temporal lobe architecture including major pathways. Tractography shows that hippocampal-thalamic connectivity was pronounced in patients with temporal lobe epilepsy [3]. Tractography is also able to detect altered hippocampus-striatum connectivity in patients with mild-traumatic brain injury and post traumatic stress disorder [10]. With diffusion tensor imaging (DTI), it has been shown that in patients with AD, changes in diffusion properties were found in the parahippocampal white matter, and in regions that are closely connected to the medial temporal lobe (MTL) [11]. DTI has also been used in patients with AD to show a significant reduction of white matter fractional anisotropy (FA) and increase in radial diffusivity in the bilateral uncinate fasciculus, parahippocampal cingulum, and fornix [12]. Diffusion parameters of the hippocampus, like mean diffusivity (MD), were also shown to be an excellent predictor of age [11].

DTI at ultrahigh sub-millimeter resolution allows us to successfully isolate the signal from the PP in the hippocampus. This approach has been used to show age-related degradation of the PP in older adults [12]. The PP has also been virtually reconstructed using DTI using ex-vivo MTL samples which has been validated with histology [13]. Diagnosis of diseases that affect the MTL like AD and mesial temporal sclerosis are particularly challenging due to the difficulties associated with identifying decisive imaging features. DTI has shown promise to alleviate these difficulties. Using near-millimeter isotropic voxel size and sophisticated acquisition schemes, along with a comprehensive segmentation of the MTL, a detailed map of the MTL architecture was made with connectivity maps from tractography closely resembling animal studies, such a methodological approach can be used in clinical settings to detect disruption of pathways due to cell loss [14]. dMRI has also been successful in mapping the macroscopic neuronal projections of the mouse hippocampus. Tractography from high spatial and angular resolution dMRI scans of the mouse brain expose the organizational pattern of dendrites and axonal networks in the hippocampus which are highly correlated with histological tracer studies [15]. These findings show the potential of dMRI and tractography for studying neuronal projections in complex brain regions like the hippocampus.

In this work, we introduce a new model which employs anatomical coordinates to virtually unfold the hippocampus into a thin sheet [16] and project vector-valued diffusion data onto this unfolded hippocampus. We demonstrate that by utilizing our method, open access 3T data from the Human Connectome Project can be used to map the complex anatomy of the hippocampus. We are able to investigate the inner connectivity of the hippocampus and recover elements of the polysynaptic pathway which are difficult to discern in the native configuration of the hippocampus in an in vivo 3T scan. As a proof of concept, we show that with moderately high resolution in vivo human MRI, our method reproduces patterns of connectivity that were shown to exist in tracer-based studies in primate.

## 2 Methods

### 2.1 Virtual unfolding of hippocampus with Laplace’s equation

Laplace’s equation, ∇^2^*ϕ* = 0, where *ϕ* is a scalar field, is ubiquitous in nature. In the realm of computational neuroimaging, Laplace’s equation has been used as a robust tool to accurately calculate the thickness of the neocortex [17], [18]. One attractive feature of a solution to Laplace’s equation is a smooth (twice-differentiable) transition of the scalar field from one boundary to another. We use this feature to find three harmonic coordinates *u, v* and *w* for the hippocampus. In Figure 1 we see the boundaries chosen for each coordinate (e.g., when solving for the coordinate *u, U*_0_ is the source and *U*_1_ is the sink). The boundaries of these coordinates are motivated neuroanatomically, and consist of structures which border the hippocampus at its topological edges. For example, the hippocampal-amygdalar transition area and induseum grisium represent natural anterior and posterior termini of the hippocampus, respectively (see [16] for additional details on how these boundaries are defined). We choose Neumann boundary conditions on the rest of the hippocampus to ensure that the tangent vectors of the coordinates are orthogonal to the surface normals at the boundary of the hippocampus. When solving for the coordinates *v* and *w*, we follow an analogous procedure (e.g., when solving for *v* we chose the source and sink as *V*_0_ and *V*_1_ respectively), and Neumann boundary conditions on the rest of the hippocampus. The solution to Laplace’s equation is obtained by the algorithm of [16] with two changes, first, the coordinates were reparameterized by arc length so that the unfolded hippocampus would still preserve its actual length dimensions in SI units. Second, the coordinates were linearly extrapolated into the hippocampal sulcus to ensure that the stratum radiatum/stratum lacunosum-moleculare (SRLM) was captured in our unfolded hippocampus. Figure 1 shows the solved coordinates.

**Figure 1:**
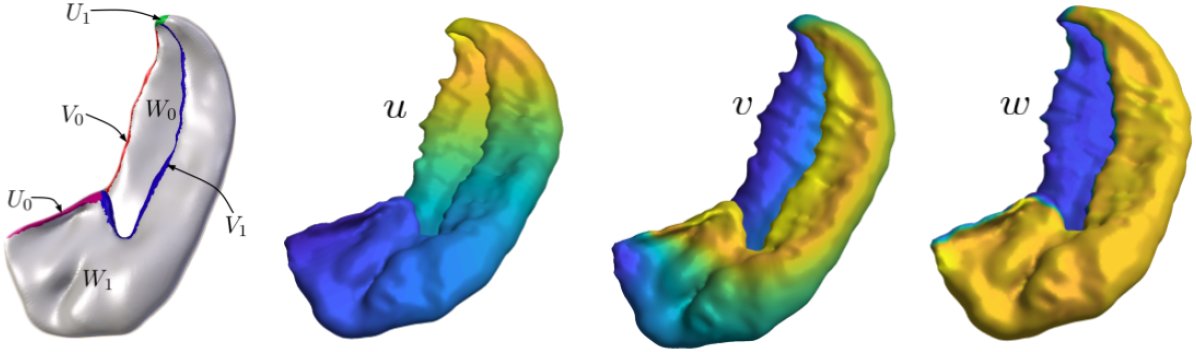
This figure shows a surface rendering of a manual segmentation from an HCP subject. The leftmost image shows the regions *U*_0_, *U*_1_, *V*_0_, *V*_1_, *W*_0_ and *W*_1_ which were manually chosen for boundary conditions for Laplace’s equation. Note that *W*_0_ is the inner surface of the hippocampus and *W*_1_ is the outer surface. The remaining images show the solutions obtained for each field *u, v* and *w*. The coordinates increase from blue to yellow.

Given an intensity *I*(*x, y, z*) and coordinate fields *u*(*x, y, z*),*v*(*x, y, z*) and *w*(*x, y, z*), we are able to construct the mapping *I*(*x, y, z*) → *I*(*u, v, w*). The image *I*(*u, v, w*) is interpreted as an unfolded hippocampus. Hereafter we refer to the *u, v, w* space as the ‘unfolded space’ and the *x, y, z* space as the ‘native space’, this is depicted in Figure 2.

**Figure 2:**
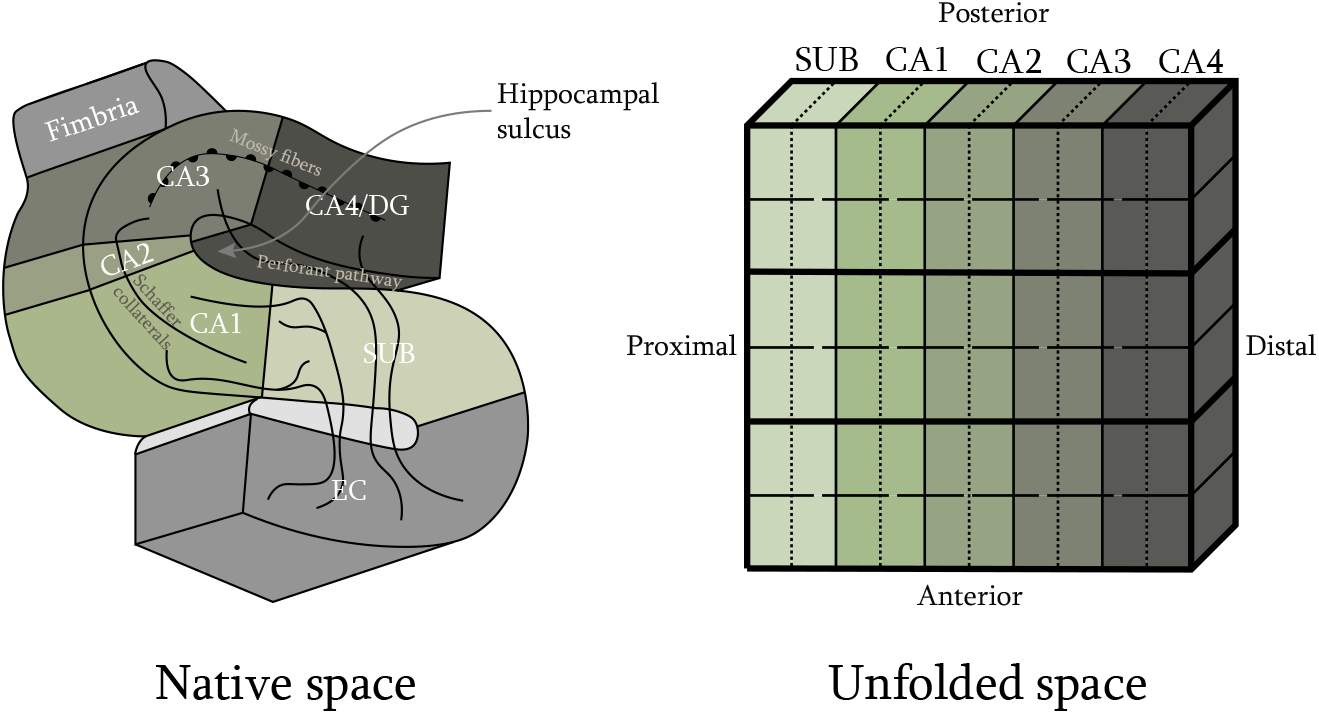
A schematic of the unfolding, on the left we have the native space with the main pathways of the hippocampus showing, on the right is the unfolded space with the subfields illustrated. Note that the EC and the fimbria are not part of our unfolding. To infer intra-hippocampal connectivity we split the hippocampus further, each subfield is split into a distal and a proximal part and the anterior-posterior length of the hippocampus is split into six regions.

### 2.2 Study data and manual segmentation

Preprocessed data was taken from the Human Connectome Project (HCP) Young Adult 3T study, WU-Minn S1200 release [19]. The dMRI scans are multi-shell with b-values of 0, 1000, 2000, and 3000 s/mm^2^ with approximately an equal number of acquisitions on each non-zero b-shell and an echo spacing of 0.78 ms. The resolution is 1.25 mm with isotropic voxels. Further details of this dataset can be found at [20]. The virtual unfolding currently requires manual segmentation of the SRLM and gray matter of the hippocampus (utilizing the 0.7mm isotropic resolution 3D T2-weighted images), thus we limited the investigations in this work to 8 hippocampi (left and right, from 4 subjects). Manual segmentation was carried out using a protocol similar to our previous work [16].

### 2.3 Creating a diffusion weighted image for the unfolded hippocampus

We saw above how a scalar intensity *I*(*x, y, z*) can be easily mapped to the unfolded space to get *I*(*u, v, w*). Diffusion data, however, is vector-valued since a gradient direction is associated with each image. Hence we need to map vectors from the native space to the unfolded space. This can be achieved with the Jacobian matrix,

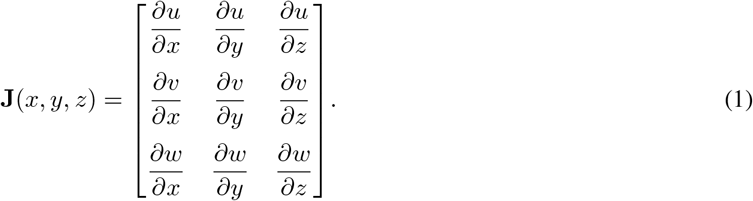

Given a vector in the native space 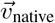, it can be transformed to the unfolded space by 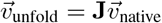. This mapping allows us to map diffusion gradient directions to the unfolded space.

A caveat of the unfolding map is the introduction of shear deformations which leads to incorrect interpretation of the diffusion acquisition resulting in overstated anisotropy values. To circumvent this issue we borrow techniques from continuum mechanics and perform a polar decomposition of the Jacobian matrix, **J** = **RU** where **R** is the rotation matrix and **U** is the deformation matrix. We take only the rotation part and set **U** = 1 to obtain **J** = **R**.

Finally, we can generate a diffusion weighted image of the unfolded hippocampus. This can be achieved in a straightforward manner by taking the intensity *I*(*x,y,z*) in native diffusion volumes and mapping them to *I*(*u,v,w*) but accompanied by a mapping of the diffusion gradient directions to the unfolded space. For each gradient direction 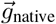, the corresponding vector in the unfolded space is given by, 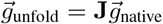. It is crucial to note here that the Jacobian matrix **J** has spatial dependence which will be carried over to the gradient directions in the unfolded space, making 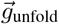 a function of space even though 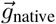 ideally does not have a spatial dependence. We generate unfolded volumes at two resolutions, 1.25 mm voxels to match the original HCP native space volumes and upscaled volumes at a resolution of 0.625 mm. For comparisons, we also upscale the native space volumes to this resolution.

### 2.4 Model fitting in the unfolded space

We fit two models to the unfolded and native space hippocampi. Diffusion tensor fitting with a weighted least squares approach (implemented in FSL as `DTIFIT` [19] and a ball-and-stick partial volume model capable of modeling multiple fibre populations (implemented in FSL as `BEDPOSTX` [20]). To account for the spatial dependence of 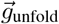 we interpret the Jacobian matrix **J**(*x,y,z*) as a gradient non-linearity. Both `DTIFIT` and `BEDPOSTX` implementations allow to correct for gradient nonlinearities via the option of including a `grad_dev.nii.gz` file, which can now also be utilized for the Jacobian matrix of the unfolding map. We perform probabilistic tractography by using the `probtrackx` command from FSL. For deterministic tractography we use the `track` command from the camino toolbox. Both `probtrackx` and `track` construct tracts from the output of `BEDPOSTX` [21].

### 2.5 Subfield segmentation and connectivity

We perform a segmentation of the hippocampus subfields through a template defined in the unfolded space which was created from manual segmentations in previous work [16]. An attractive feature of our coordinates is that because their boundaries are neuroanatomically chosen the template for the subfields falls in the correct place when we solve for the coordinates *u, v, w*. This also implies that we have segmentations in both the native and unfolded spaces given that we can map data from one space to the other. Once the subfields are segmented we can use the results of our ball and stick model to perform probabilistic tractography to study the intra and inter subfield connectivity of the hippocampus. We use `probtrackx` to perform this analysis [20]. Briefly stated, `probtrackx` will emanate tracts from a seed voxel in all directions and then record for each of the other voxels how many tracts pass through it. The parameters chosen for `probtrackx` are: number of samples were taken to be 2560/mm^3^, this is chosen from the default value of 5000 samples for a voxel volume of 1.25 mm^3^, step length (the length of each line segment for the tract) was taken to be 25% of the length of a voxel which is slightly reduced from the default of 40% for smoother tracts, the curvature threshold for the tracts was set to the default value of approximately 80 degrees and the distance threshold was 0 mm which is the default value. Every voxel was taken as a seed giving a large seed to seed connectivity matrix, 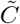, with size *N × N* where *N* is the number of voxels in the image.

For a pair of voxels *i* and *j*, the matrix entry 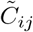 is the number of tracts passing through voxel that started in voxel *i* and conversely the matrix entry 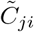 is the number of tracts passing through voxel *i* that started in voxel *j*. Hence we expect that 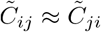. We can make this matrix smaller by collapsing the rows and columns by an average connectivity based on subfield membership of each voxel. We call this smaller matrix, *C_lm_*, which has size 5 × 5 since there are five regions in our segmentation.

When performing probabilistic tractography we can also check which trajectories are taken by tracts when they connect two subfields. If two subfields *l* and *m* are connected we can determine how many tracts go through the remaining subfields enroute from l to m or vice-versa. Using this feature we can exclude tracts that visit other subfields and only keep the ones that connect two subfields directly. For these tracts we call the resulting connectivity matrix 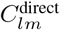. This distinction is motivated by the connectivity of the hippocampus, since we know from histological studies that the polysynaptic pathway sequentially connects subfields, only neighbouring subfields should be directly connected for a plausible result.

To study the intra-hippocampal connectivity we split the hippocampus as depicted by the dotted lines in Figure 2. We split each subfield into two regions, a proximal one and a distal one. This split is done along a surface that is midway between the distal and proximal boundaries that define the subfield. We also split the unfolded hippocampus into six regions in the anterior to posterior direction. These regions, along with the connectivity matrix 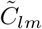, allow us to examine the topographic organization of intra-hippocampal connections. Similar to the method of investigating the connectivity of the subfields described above, we can reduce 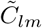 to give us connections between specific regions.

To examine the topographic organization of the connections in the hippocampus we make comparisons with findings from discrete anterograde and retrograde tracer injections in the macaque monkey hippocampal formation [22]. These tracer injections revealed certain trends in the non-human primate hippocampus that can be readily tested by our unfolding technique. For example, one trend found in the macaque hippocampus is that CA3 associational projections are less pronounced than CA3 projections to CA1. To check this using our method, we can filter the connectivity matrix 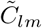 to show the strength of associational projections in CA3 and then compare that with 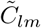 filtered to show the strength of connections between CA3 and CA1. We check the following connectivity trends which are also illustrated in Figure 3:

1. *CA3 associational projections and CA3 projections to CA1:* One trend that was revealed in tracer studies is that in the nonhuman primate hippocampus CA3 associational projections, were less pronounced than those from CA1 to CA3.
2. *CA1 and CA3 associational projections:* Another trend is that CA3 associational projections are more substantial than CA1 associational projections.
3. *CA1 associational projections and CA3 projections to CA1:* CA1 associational projections are less substantial than CA3 projections to CA1.
4. *CA3 projections to CA1 in head and tail:* CA3 projections to CA1 are more substantial in the head as compared to CA3 projections to CA1 in the tail.
5. *CA3 associational projections in head and tail:* Finally, in the nonhuman primate hippocampus CA3 associational projections in head were more substantial than CA3 associational projections in the tail.

Recently, there is also interest in studying connectivity and composition gradients along the anterior-posterior axis of the hippocampus [23], [24], [25]. Given our splitting in the anterior posterior direction we can readily investigate this with a high-level of granularity, while still respecting the curvature of the hippocampus. The unfolded hippocampus is split into six bins, 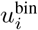 for *i* ∈ {1 … 6}. After performing probabilistic tractography we get six matrices, 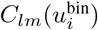, and six other matrices, 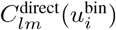.

**Figure 3:**
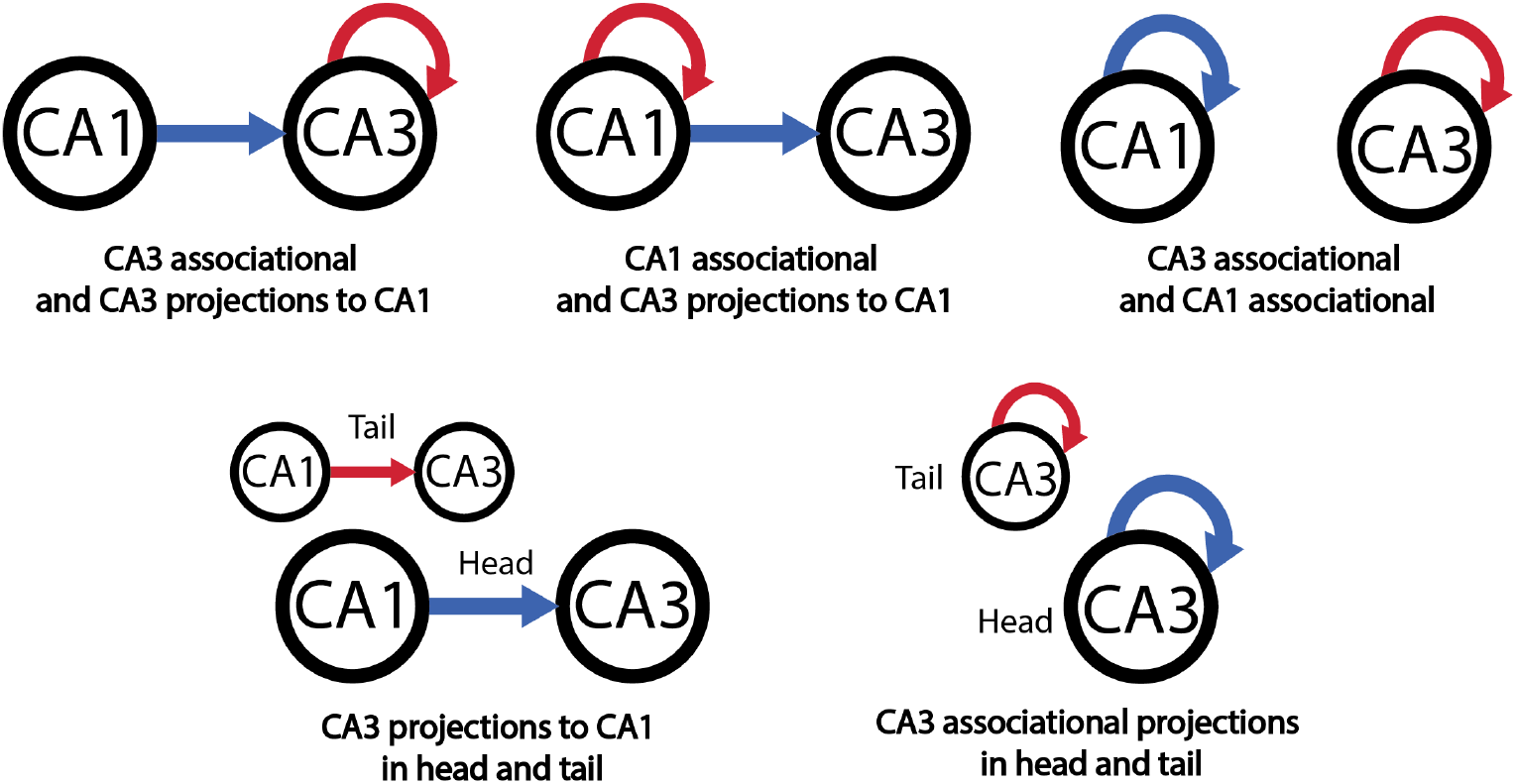
An illustration of the connectivity trends that we test with our unfolding technique. Circular arrows represent associational projections. Projections represented with blue arrows are compared with projections shown with red arrows.

### 2.6 Test of unfolding method

We perform two tests to check if our method is corrupting the diffusion data when we perform the mapping from the native space to the unfolded space. For the first test, let 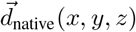 be the dyads from the ball and stick model in the native space and let 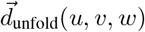 be the dyads from the unfolded space. We may then look at the dot product 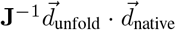 and from that calculate the angle, *θ*(*x, y, z*), between 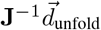 and 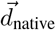. This angle should be small. Another simple test is to compare the histogram of the FA values obtained from the native space with the histogram of FA values obtained from the unfolded space. This will reveal any shearing artifacts introduced by the mapping.

## 3 Results

### 3.1 Unfolded space visualization of diffusion parameters

In Figure 4 we find the native space T2-weighted images cropped around the left hemisphere hippocampus of a participant. Overlaid on these images are the first two dyads obtained from `BEDPOSTX`. The green dyads indicate a direction from anterior to posterior, red indicates right to left and blue is inferior to posterior. In Figure 5 we see the unfolded hippocampus volume at a resolution of 0.625 mm. The background intensity is FA and overlaid are the dyads obtained from `BEDPOSTX`. The three images are slices of the hippocampus in the w or superficial to deep coordinate. Here, the top most slice is the most superficial layer and the bottom most slice is the deepest layer. In this space, red indicates anterior to posterior, green proximal to distal and blue in/out in the laminar direction.

**Figure 4:**
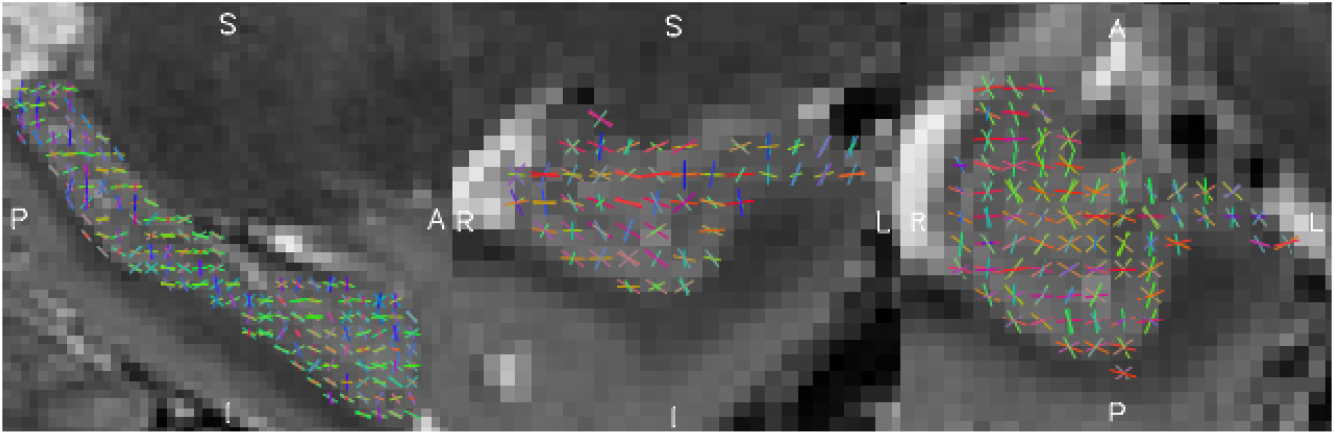
From left to right we see the sagittal, coronal and axial slices of a T2 weighted image cropped around the left hippocampus of a subject. The colored sticks show fiber orientations resulting from a `BEDPOSTX` fit at a resolution of 1.25 mm, green represents the anterior to posterior direction, red represents right to left and blue inferior to superior.

**Figure 5:**
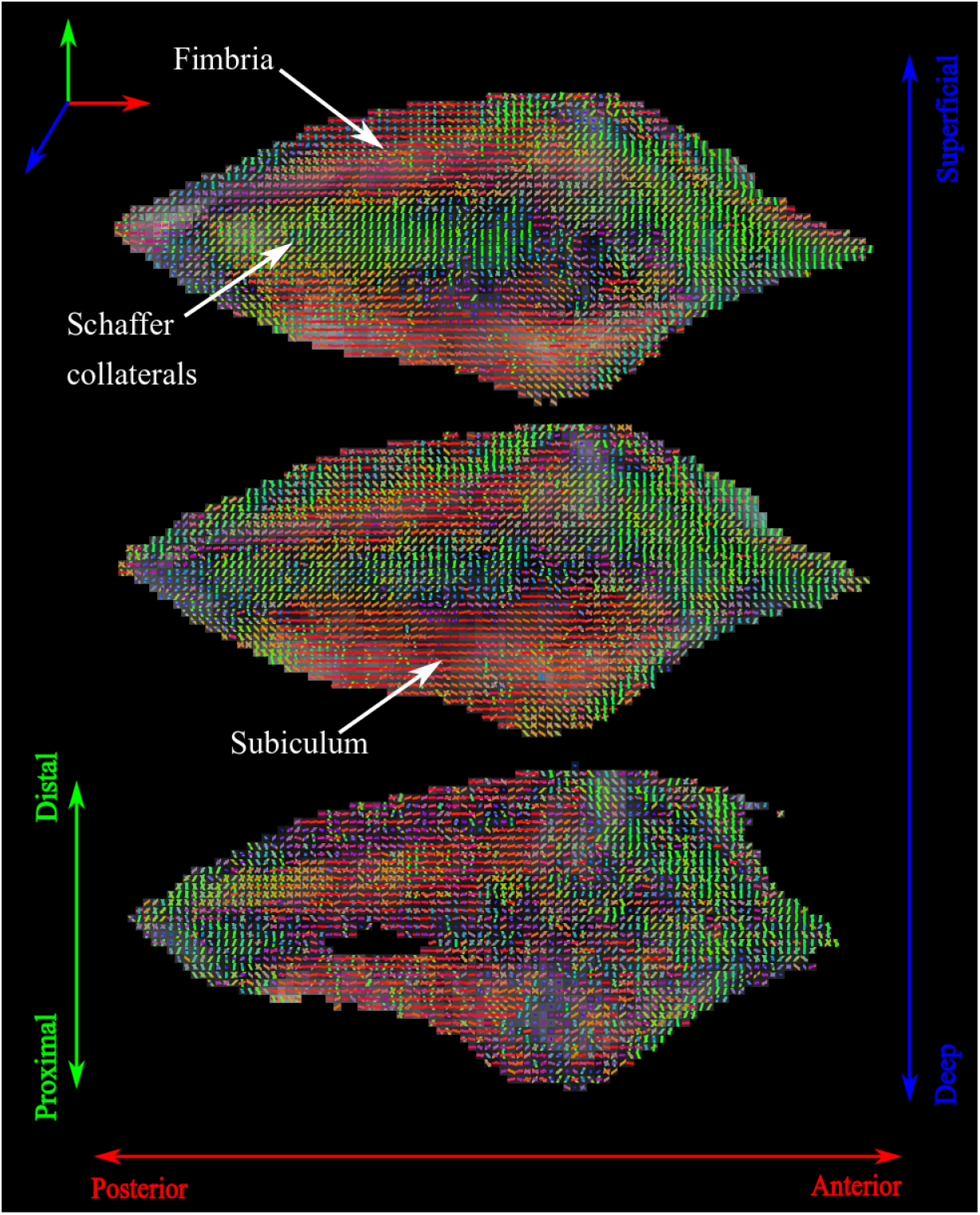
This figure shows the unfolded hippocampus at a resolution of 0.625 mm. From top to bottom we move from the most superficial surface of the hippocampus to the deepest (shown with the blue axis). For each individual slice, from top to bottom we go from distal to proximal (as shown by the green axis) and from left to right we go from posterior to anterior (shown by the red axis). The colored sticks show fiber orientations from the `BEDPOSTX` fit, green shows proximal to distal, red shows posterior to anterior and blue shows superficial to deep. Some patterns in the sticks are visible and the likely underlying structures are labelled. In the most superficial slice (top), in the distal region, we see a coherent structure going from posterior to anterior which is where we would find the fimbria. Adjacent to the fimbria we see fibres running proximal to distal, a pattern which continues in the deeper layers also, this is where one would find the Schaffer collaterals. We also find a coherent structure running posterior to anterior in the proximal part of the slices due to white matter fibres in the subiculum.

In previous work [16] we had a square unfolded hippocampus, here we see a more distorted shape. This is due to the reparametrizing of the coordinates with arc length as the new parameter which makes relative length scales more apparent. For example, hippocampi are longer in the anterior to posterior direction as compared to the proximal to distal direction; this is now illuminated in the unfolded hippocampus. We see immediately that with the unfolded space we are able to view the dyads in Figure 4 in a more convenient setting. Certain patterns are illuminated by the dyads in the unfolded space. In the most superficial layer of the hippocampus (top), in the distal part, we see a clear coherent structure running from posterior to anterior. This structure is likely due to the fimbria, depicted in Figure 2. Moving in the proximal direction from the fimbria we see coherent fibres running proximal to distal, a pattern which continues in the other slices also. Anatomically the underlying pathway is the Schaffer collaterals which also runs proximal to distal. In the middle and top layers, in the proximal region, we see another coherent structure running form posterior to anterior, this corresponds to the subiculum.

### 3.2 Test of unfolding method

As mentioned above we perform tests to verify that the implementation of our map to the unfolded space does not corrupt the diffusion data. In Figure 6a we see a histogram of the values *θ*(*x, y, z*) which is the angle between the dyads in the native space and the unfolded space, notice that the values are distributed around 0 suggesting that our method works well. Further note that the angles are between 0 and 90, this is because dyads are directionless and for a *θ* > 90 we take the angle as 180 - *θ*. Figure 6b shows two normalized histograms, the orange bars in the foreground are for FA values obtained in the native space and the blue bars in the background are for FA values in the unfolded space. Notice how the distribution in both spaces is close to identical, this agreement is a direct result of our approach of setting the shear part of our transformation to the identity map.

**Figure 6:**
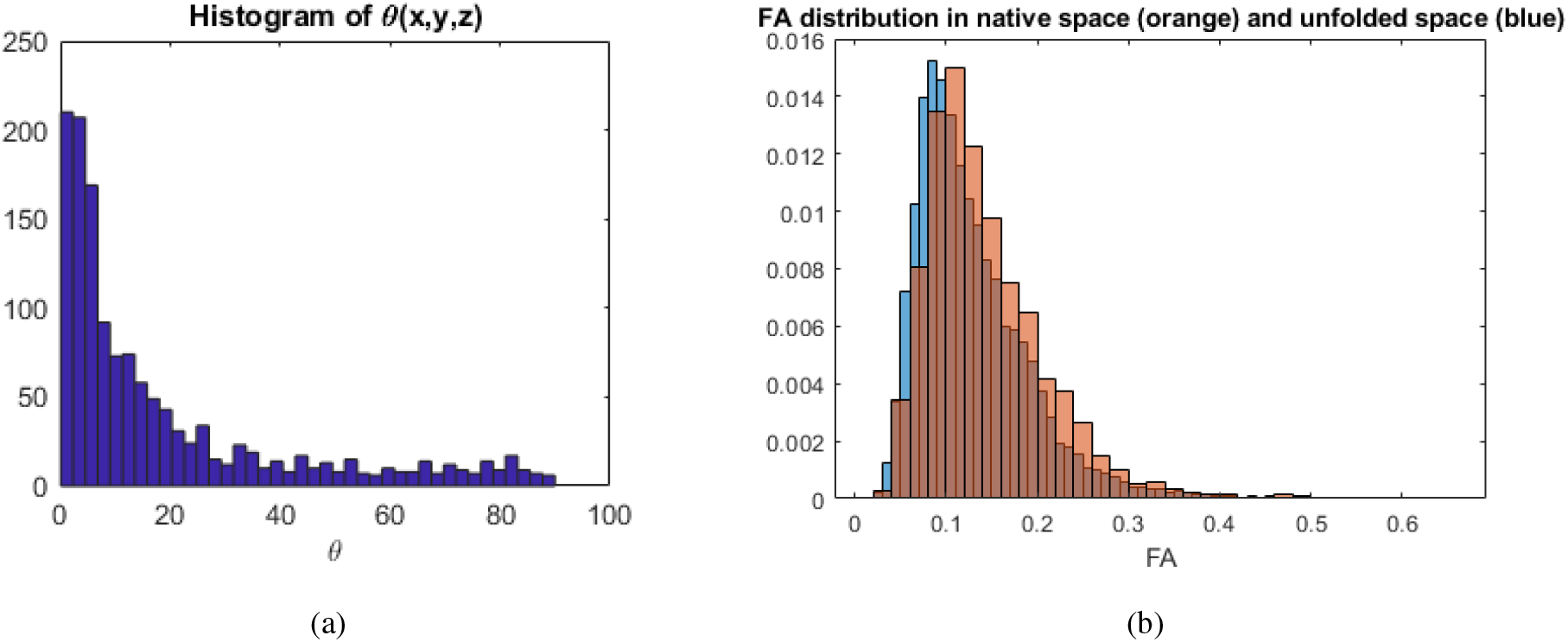
a) shows *θ*(*x, y, z*) which is the angle between the dyads in the native space and the unfolded space. Here, the dyads in the unfolded space have been moved to the native space by using the inverse of the Jacobian matrix. b) shows two normalized histograms for FA values on the hippocampus. The blue histogram in the background is for FA values in the unfolded space and the orange in the foreground is for FA values in the native space. Notice how they are close to identical, validating our method.

### 3.3 Subfield segmentation

A benefit of our unfolded space is that defining the boundaries of the subfields can be performed by employing the *u, v, w* coordinates directly, and does not rely on heuristics based on the appearance of the hippocampus in a standard coronal section. In this way, we are able to perform a reliable segmentation of the subfields throughout the extent of the hippocampus, as demonstrated in Figure 7. This segmentation is done with a template (determined from our previous work [16]) which is defined on the the *u, v, w* coordinates, and thus automatically registered once we solve for these coordinates.

**Figure 7:**
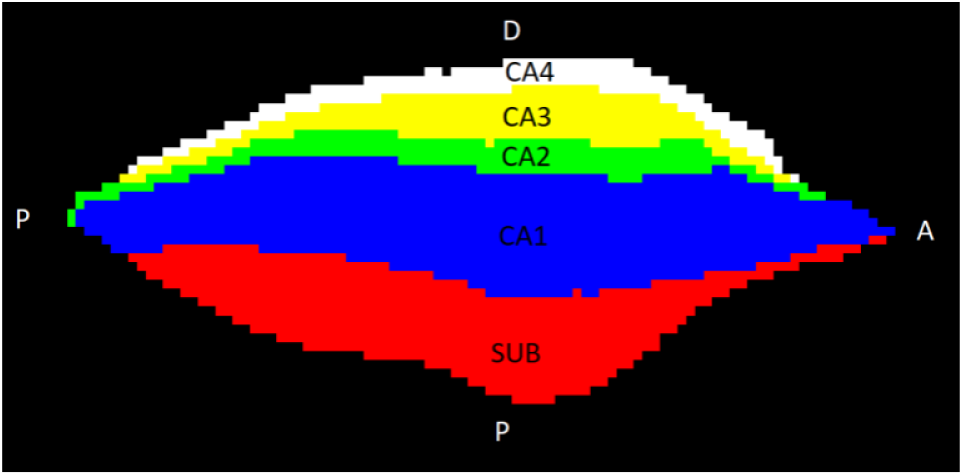
An example of the automatic subfield segmentation from templates defined on the anatomical coordinates.The top most region in white is the dentate gyrus or CA4, followed sequentially by CA3-CA1 (yellow, green and blue) and then ending with the subiculum in red.

### 3.4 Connectivity

Figure 8 summarizes our findings from probabilistic tractography performed at a resolution of 0.625 mm, it shows for both the native and unfolded space the mean and standard deviation over all subjects for the matrices *C_lm_* (Indirect) and 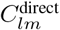 (Direct). The matrices for each participant are shown in the appendix. The off diagonal components show the inter-connectivity between two subfields and the diagonal components are the intra-connectivity. In the unfolded space, we see that both matrices, *C_lm_* and 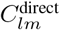, are more diagonal, in the sense that entries beyond the super-diagonal and sub-diagonal are smaller (the super-diagonal and sub-diagonal are shown in green in Figure 9). This signifies that subfields that are adjacent are the most connected, which is a result in tune with the anatomy of the hippocampus. This effect is summarized in the table in Figure 9, where the entries are the ratio of the sum of the green entries, to that of the sum of yellow entries. Here we can clearly see that in the unfolded space adjacent subfields are highly connected. In Figure 10 we see the results of probabilistic tractography after performing an anterior to posterior binning. Figure 10 a) shows the diagonal entries, 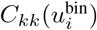, b) shows the super-diagonal, 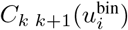. Figure 10 c) and d) are analogous to a) and b) respectively with the exception that we use direct tracts, i.e, matrices 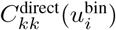 and 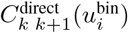.

**Figure 8:**
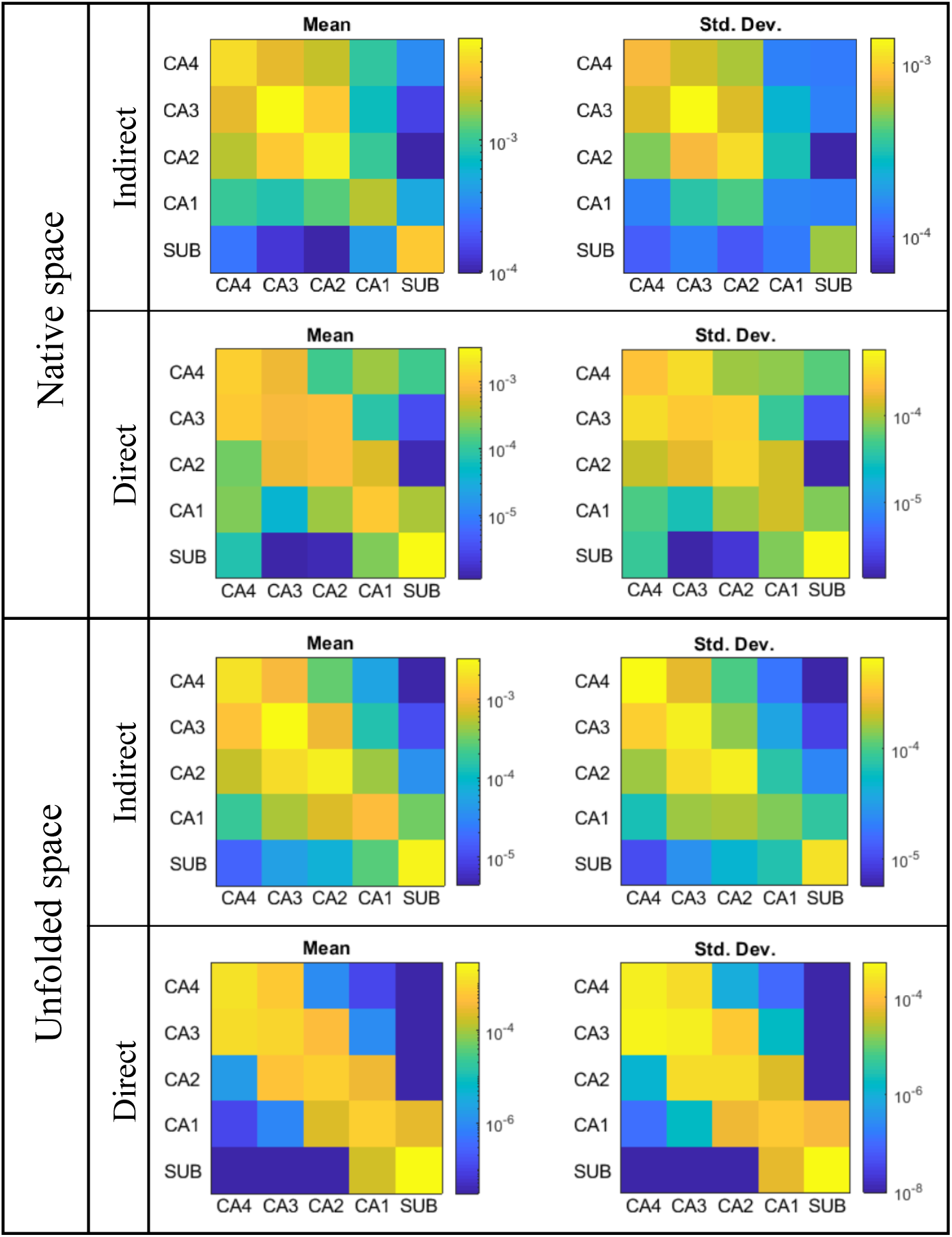
Here we see the mean and standard deviation of connectivity matrices generated from probabilistic tractography, performed at a resolution of 0.625 mm, for both the native space and the unfolded space. For each of the spaces we have the indirect connectivity matrix, *C_lm_*, and the direct connectivity matrix, 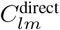. We see that *C_lm_* for the unfolded space is more anatomically plausible as there are fewer connections going beyond the super and sub diagonals as compared to the native space. Further, from 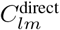 this is more apparent, we see clearly that in the unfolded space subfields are high connected to only adjacent subfields. In the native space, 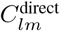 reveals that there are spurious, implausible connections connecting non-adjacent subfields.

**Figure 9:**
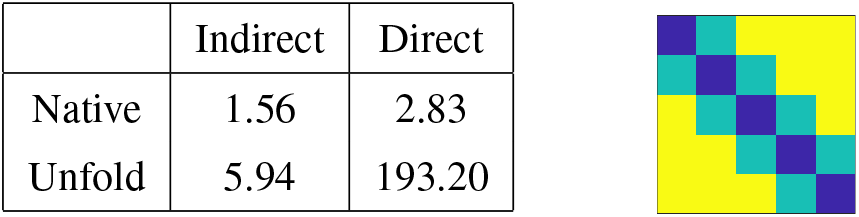
In the table we see the ratio of the sum of the values in the green regions to the sum of the values in the yellow region. This ratio estimates how connected subfields are to their immediate neighbours versus distant neighbors. We can clearly see that in the native space this ratio is low as compared to the unfolded space, this implies that in the unfolded space we get more connections to immediate neighbors of subfields which is anatomically more plausible.

**Figure 10:**
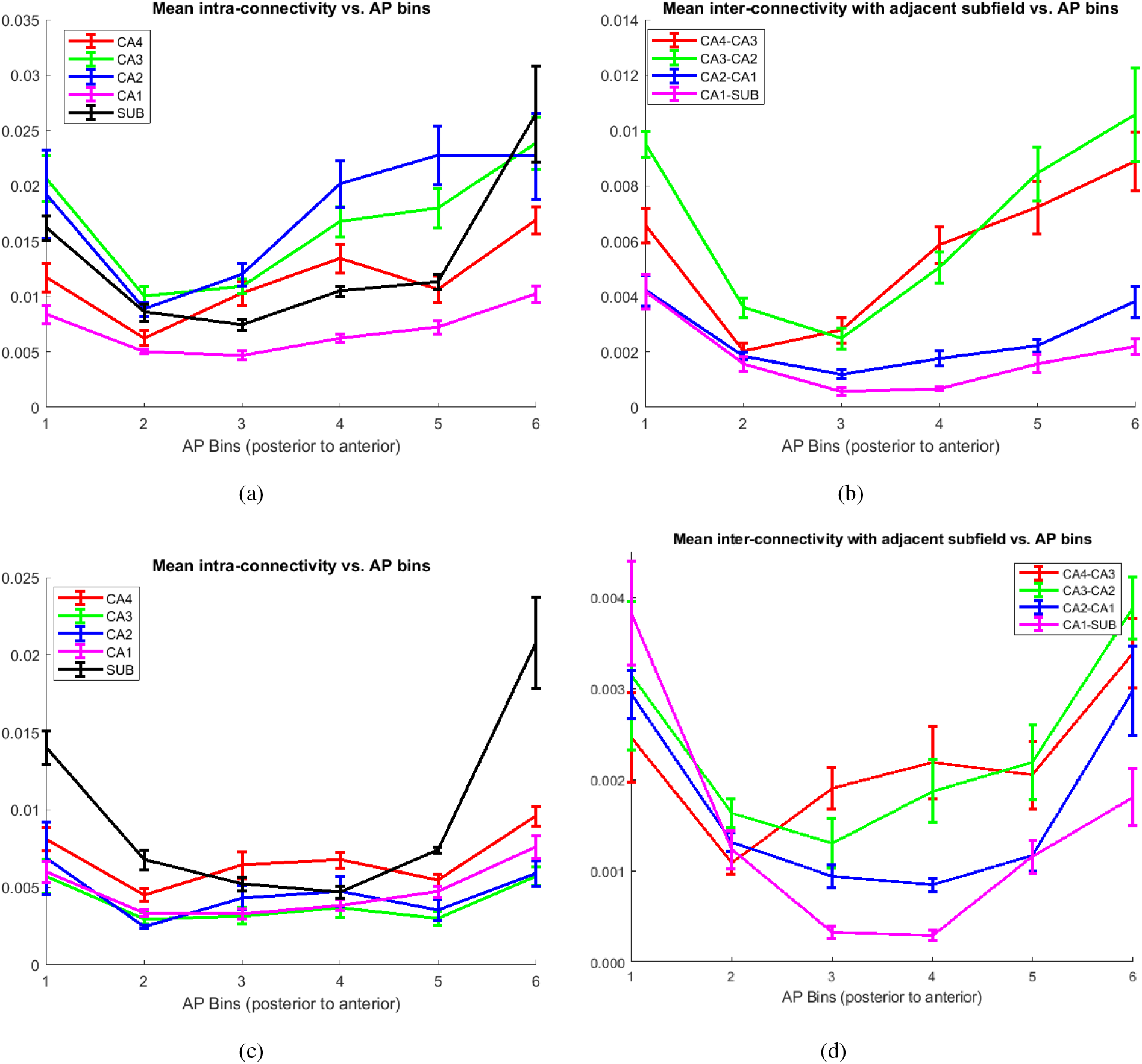
In a) we see the average indirect intra-connectivity in the unfolded hippocampus as function of anterior to posterior bins. In b) we see the average indirect interconnectivity in the unfolded hippocampus as a function of anterior to posterior bins. c) and d) are analogous to a) and b) respectively with the exception that the tractography tracts are direct. We notice, in general, a trend of higher intra and inter connectivity in the anterior (head) of the hippocampus. The error bars are the standard error.

As described above, to examine the intra-hippocampal connections we look for the existence of certain trends that have been previously observed from histological studies using tracer injections in the nonhuman primate hippocampus [22]. The resulting connectivity values are summarised in Table 1, also shown are the p-values from a two-sample t-test. The connectivity values are averaged over all hippocampi and we use the indirect connectivity matrix, 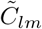, calculated in the unfolded space at a resolution of 0.625 mm. Note that, except for one entry, all our connectivities follow the trends from the tracer studies with a p-value of less than 0.05.

**Table 1:**
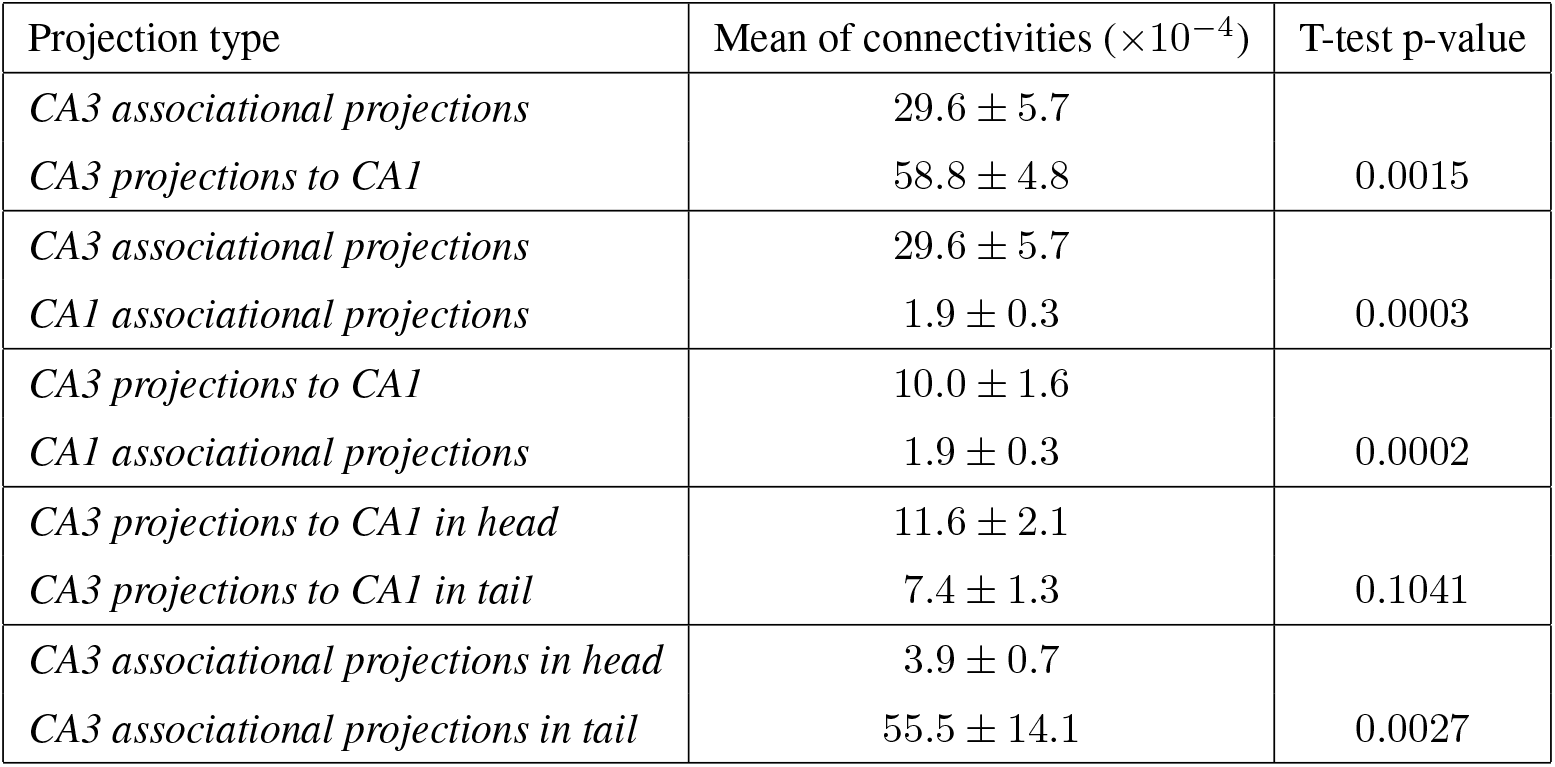
This table summarizes our findings for intra-hippocampal connectivities. For each projection type we see the mean connectivity with standard error. Also shown are the p-values from performing a two-sample t-test. With the exception of CA3 projections in the head and tail we are able to verify all the connectivity trends with a p-value of less than 0.005. Although the p-value for CA3 projections in the head and tail is high, the trend is correct, i.e., projections in the head are more pronounced than in the tail.

### 3.5 Deterministic tractography

We use the track command from the camino toolbox to perform deterministic tractography at a resolution of 0.625 mm. These tracts are shown in Figure 11 and Figure 12. In Figure 11 we see the top down and bottom up view of a pair of hippocampi. Of the pair, the ‘Native’ tracts are generated by performing tractography in the native space and the ‘Unfolded’ tracts are generated in the unfolded space and then mapped back to the native space.

**Figure 11:**
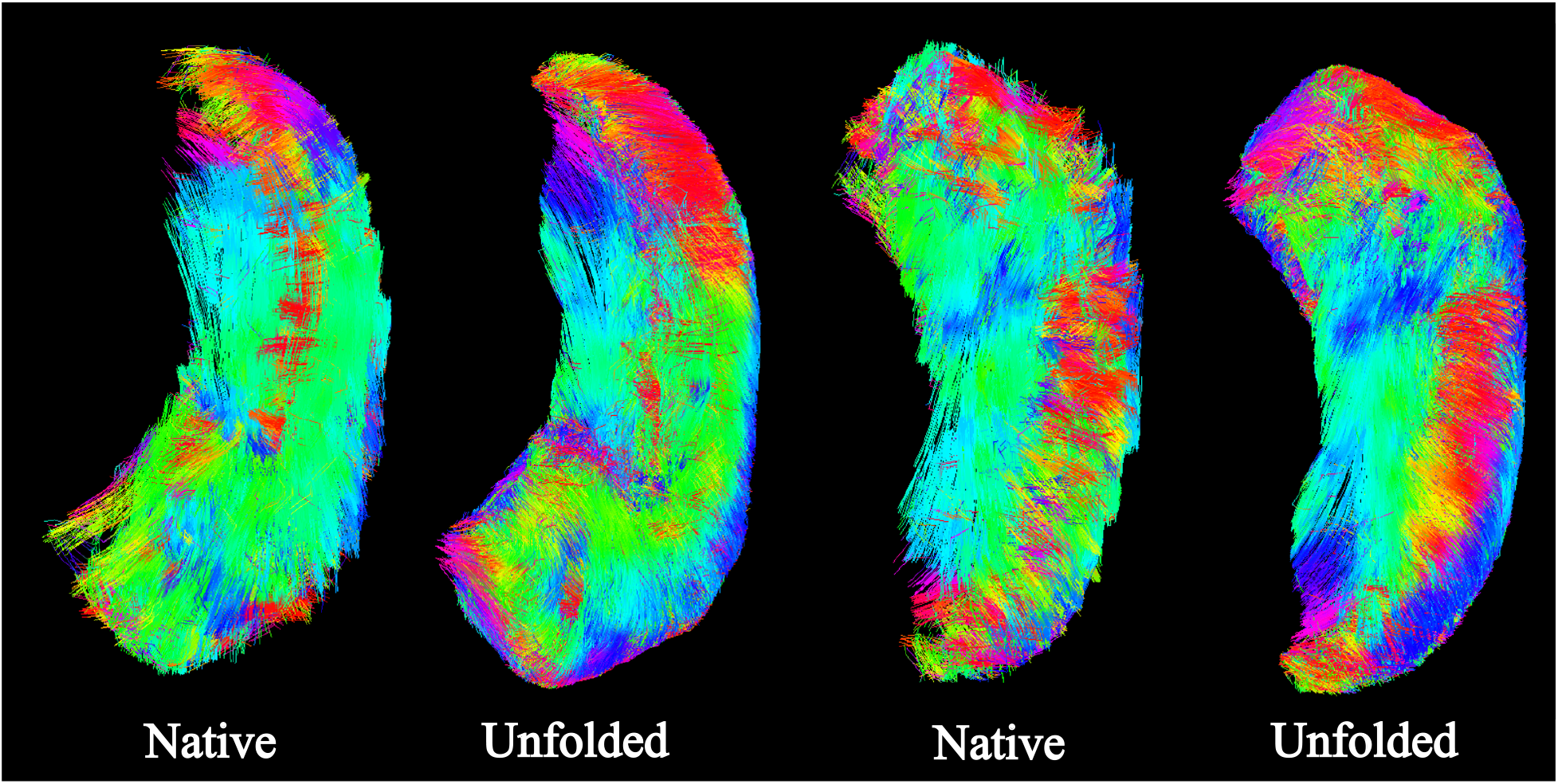
These images show the results of deterministic tractography at a resolution of 0.625 mm realized in the native space. The left pair of hippocampi is the top down view and the right pair is the bottom up view. Of the pairs, the ‘Native’ one is with tracts generated by performing tractography in the native space. The ‘Unfolded’ one is with tracts generated in the unfolded space and then mapped to the native space. The color coding of the tracts is the same as Figure 4.

**Figure 12:**
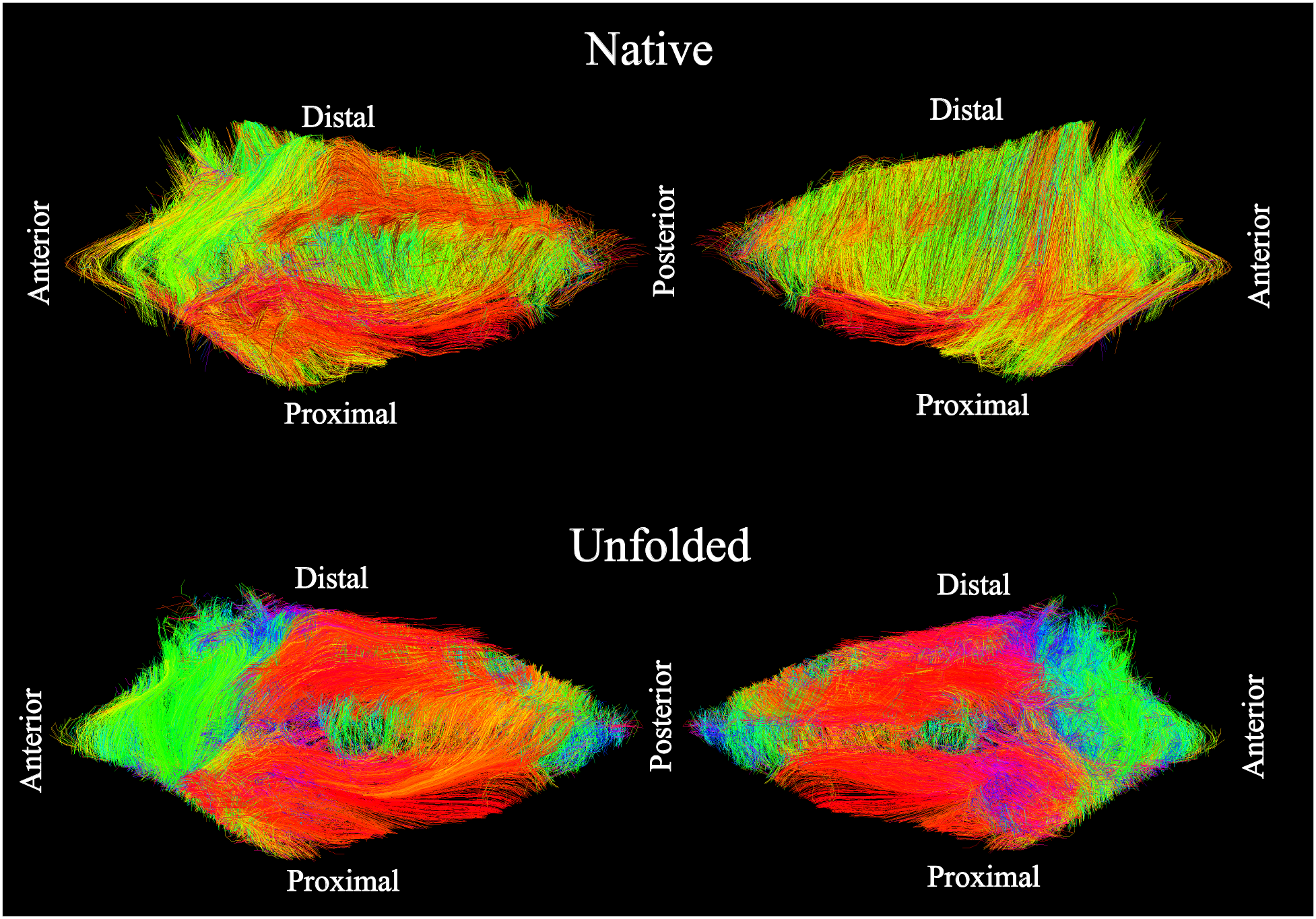
These images show the results of deterministic tractography at a resolution of 0.625 mm realized in the unfolded space. The left-top and left-bottom images are such that the most superficial slice is visible, the right-top and right-bottom are such that the deepest slice is visible. The top images are made with tracts generated in the native space and then mapped to the unfolded space. The bottom images are made from tracts generated in the unfolded space. The color coding of the tracts is the same as Figure 5.

In Figure 12 we again see two pairs of unfolded hippocampi. The top-left and bottom-left view is such that the most superficial slice is visible. Similar to the native space the top tracts are mapped to the unfolded after performing tractography in the native space and the bottom ones are generated from performing tractography in the unfolded space. The top-right and bottom-right are analogous except the view is such that the deepest slice is visible. The color coding in Figure 11 is same as in Figure 4 and the color coding in Figure 12 is the same as in Figure 5.

## 4 Discussion

In this study we have presented an approach for analyzing dMRI data in areas of the brain with complex geometries. By using an anatomical coordinate system we can flatten a convoluted region of the brain and then map the vector valued diffusion data to this flattened region. We applied this modelling approach to the hippocampus which has a complicated and curved shape. We chose anatomical harmonic coordinates with boundary conditions specified in a manner such that the axes of the coordinates now point along major pathways of the hippocampus. This allowed us to enhance our interpretation of the diffusion signal from the hippocampus. Below we discuss the key aspects of this new modelling approach, limitations of our work, and a conclusion.

### 4.1 Connectivity of the hippocampus

In the literature concerning the connectivity of the hippocampus the polysynaptic pathway is agreed upon as a standard connectiontional view [8]. One of the most prominent features of this pathway is that it connects the subfields in a sequential manner. Starting from CA4 and the dentate gyrus (DA) we see fibers going into CA2/CA3, called the Mossy fibers. From there we have the Schaffer collaterals going into CA1, and CA1 projecting onto the subiculum. This kind of sequential connectivity would translate to a connectivity matrix where *C*_*i i*+1_ and *C*_*i i*−1_ would show a large number of tracts passing through, that started in subfield i, as compared to the other subfields which should show a relatively less number of fibers passing through. Or in other words, the super-diagonal and sub-diagonals would have the highest connectivity relative to the other entries in the connectivity matrix (Figure 9). When we perform an analysis of the connectivity of the hippocampus in its native configuration, without using our new techniques, we get a result that is not in good agreement with the polysynaptic pathway. In Figure 8 (Native space → Indirect) we can see from the mean of *C_lm_* that the inter-connectivity of topologically non-adjacent subfields is overstated. This is most prominent for CA4 where we see that it has connections going to all the subfields, even the subiculum. Whereas for the native space in Figure 8 (Native space → Indirect), we obtain a result that is in agreement with the polysynaptic pathway and spurious connections do not contaminate the results. Figure 8 also shows 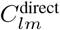, for the native space (Native → Direct) there are implausible tracts that directly connect subfields which are not sequential. Again, here we can see that there are direct paths from CA4/DG to all the other subfields. This is due to incorrect connections from CA4/DG going through the hippocampal sulcus (shown in Figure 2) to all the other subfields. With 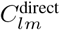 calculated by using our model, in Figure 8 (Unfolded space → Direct) we can see that there are no spurious connections going through the hippocampal sulcus to the other subfields and only neighboring subfields are directly connected. This is a direct consequence of unfolding the hippocampus as now the CA4/DG subfield is at a terminal boundary and the only region next to it is CA3 (Figure 2). We also gain more homogeneity over our participants when using the unfolded space, this can be seen in the individual participant results in the appendix. In the end we get results that are more anatomically correct and are in accordance with the polysynaptic pathway.

### 4.2 Intrahippocampal connectivity

Tracer injection studies of the non-human primate hippocampal formation revealed certain connectivity trends that we analysed above using our unfolding technique [22]. Four out of the five comparisons of the mean connectivities were in agreement with the tracer studies, all these four mean comparisons had a p-value of less than 0.05 as calculated for a 2-sample t-test. The exception to this was the trend that CA3 projections to CA1 are more substantial in the head of the hippocampus than in the tail, although comparisons of the means show the correct trend, the p-value is greater than 0.05. This lack is likely due to the limited resolution in the tail of the hippocampus. Here we also note the utility of mapping diffusion data to the unfolded hippocampus, as such an analysis would be difficult to perform in the native configuration. There are connectivity trends reported in [22] that involve the laminar depth of the hippocampus. Given the limited resolution of 1.25mm/voxel, the laminar depth of the hippocampus suffers from partial voluming artifacts and such tests are unreliable and out of the scope of this study.

### 4.3 Anterior to posterior connectivity

In Figure 10 we see the results of the anterior to posterior binning to investigate connectivity gradients. We see a general pattern where there is an increase in both the inter-connectivity, and the intra-connectivity with adjacent subfields, as we move towards the head of the hippocampus. This is a trend that has been observed in ex-vivo diffusion studies of the hippocampus also [26]. Notably, anatomical partitioning of the hippocampus from posterior to anterior in such a manner would be difficult in the native space as the underlying grid does not align with the anterior to posterior length of the hippocampus.

### 4.4 Deterministic tractography

In Figure 11 and Figure 12 we see the results of deterministic tractography. In Figure 11 the tracts are realized in the native space. Here, we see that the tracts calculated in the unfolded space and then mapped to the native space are smoother. This is because, in the unfolded space, the curvature that the tracts encounter is much lower as compared to the native space. Further, there is interpolation of the diffusion data when mapping to the unfolded space. This interpolation is performed using the anatomical coordinates rather than the cartesian grid which translates into smoother tracts. In Figure 11 we also see, visually, that the general orientation of the tracts is similar, i.e, our approach is not introducing any implausible artifacts.

### 4.5 Limitations

Currently, a limitation of our work is that it relies on a manual segmentation of the hippocampus. However semiautomatic methods for segmentation are currently in development which will help to significantly improve sample size and offer more insight into the connectivity of the hippocampus. Despite this limitation in sample size, we are still able to demonstrate the utility of the unfolding technique for tractography, by reproducing findings from primate tract tracing and ex vivo dMRI studies. Finally, given that our domain is limited to the hippocampus we are unable to model tracts that are not fully bound by such a domain, like the perforant pathway.

## 5 Conclusion

We developed a new approach to model dMRI data of complicated brain regions which involved flattening the region by the use of appropriate anatomical harmonic coordinates. We then applied this model to the hippocampus from 3T dMRI scans and studied their inter and intra connectivity. We demonstrated that even at a limited resolution of 1.25 mm, by using our novel approach, we were able to recover key elements of the polysynaptic pathway which were not recoverable in the native space due to spurious tracts passing through the hippocampal sulcus. Using the flattened hippocampus we were also able to show a connectivity gradient, such that the head of the hippocampus is more connected than the tail. Further, our method revealed connectivity trends that were found in tracer studies of non-human primates. Finally, we also showed improvements in tractography by using our method. All these enhancements stem from the implicit incorporation of hippocampal macro-geometry into the tractography fitting algorithms. In future work we aim to perform our analysis with a larger set of subjects by employing a semi-automatic hippocampus segmentation algorithm. In addition, the model developed here is general and can likely be used to analyze the neocortex.

## 6 Appendix

Following are the connectivity matrices of the individual participants which were used to calculate the mean and standard deviation in Figure 8.

**Figure 13:**
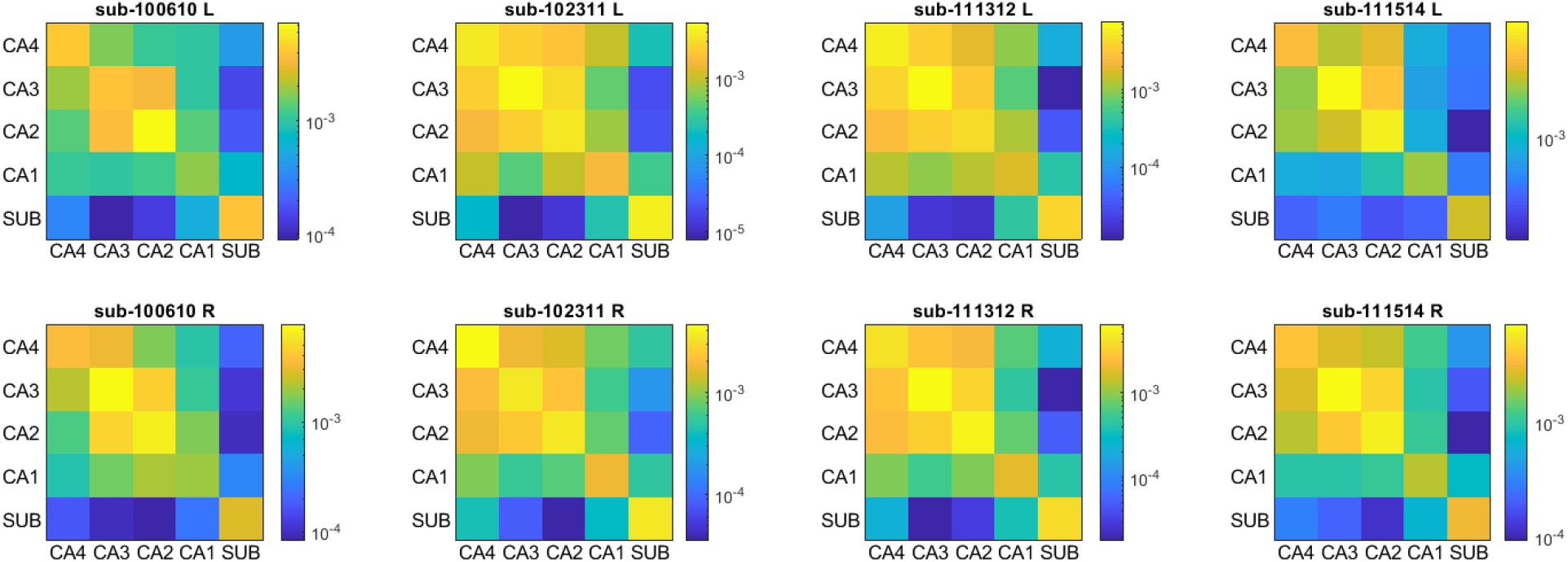
This figure shows the connectivity matrices *C_lm_* as calculated from `probtrackx` for the native space at a resolution of 0.625 mm. The columns are the different participants and the top and bottom rows are the left and right hemispheres respectively.

**Figure 14:**
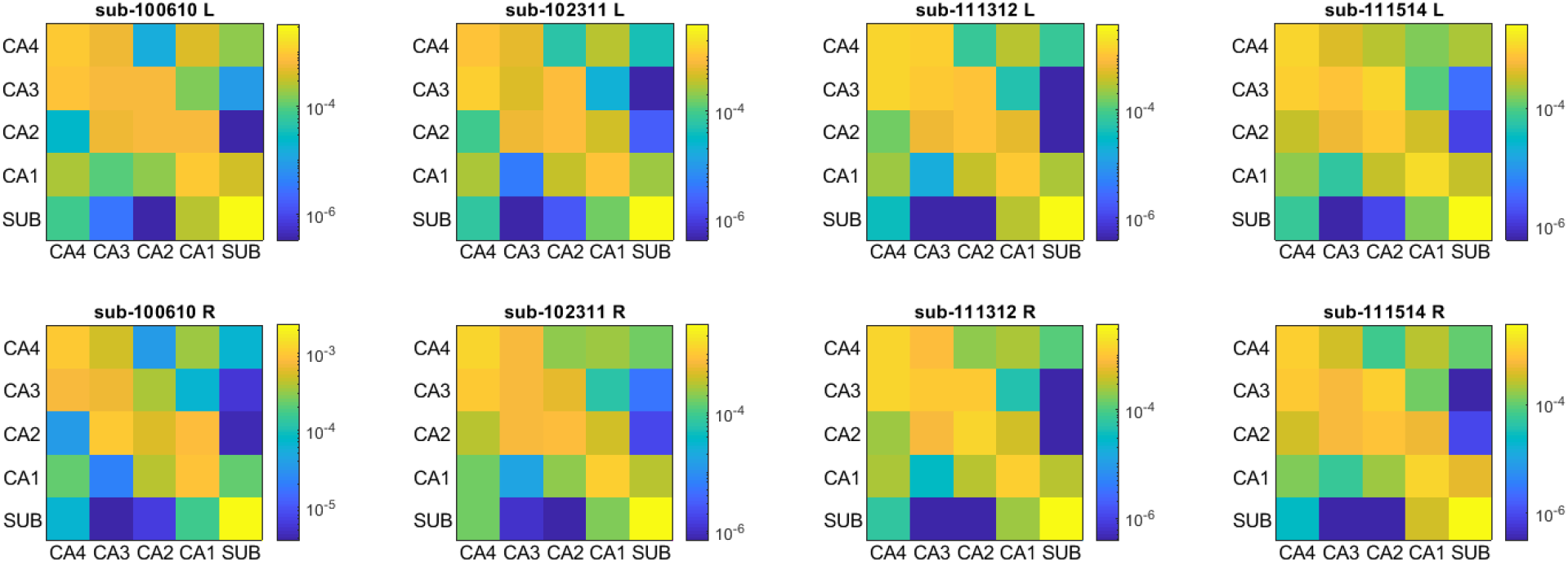
This figure shows the connectivity matrices 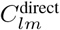 as calculated from `probtrackx` for the native space at a resolution of 0.625 mm. The columns are the different participants and the top and bottom rows are the left and right hemispheres respectively.

**Figure 15:**
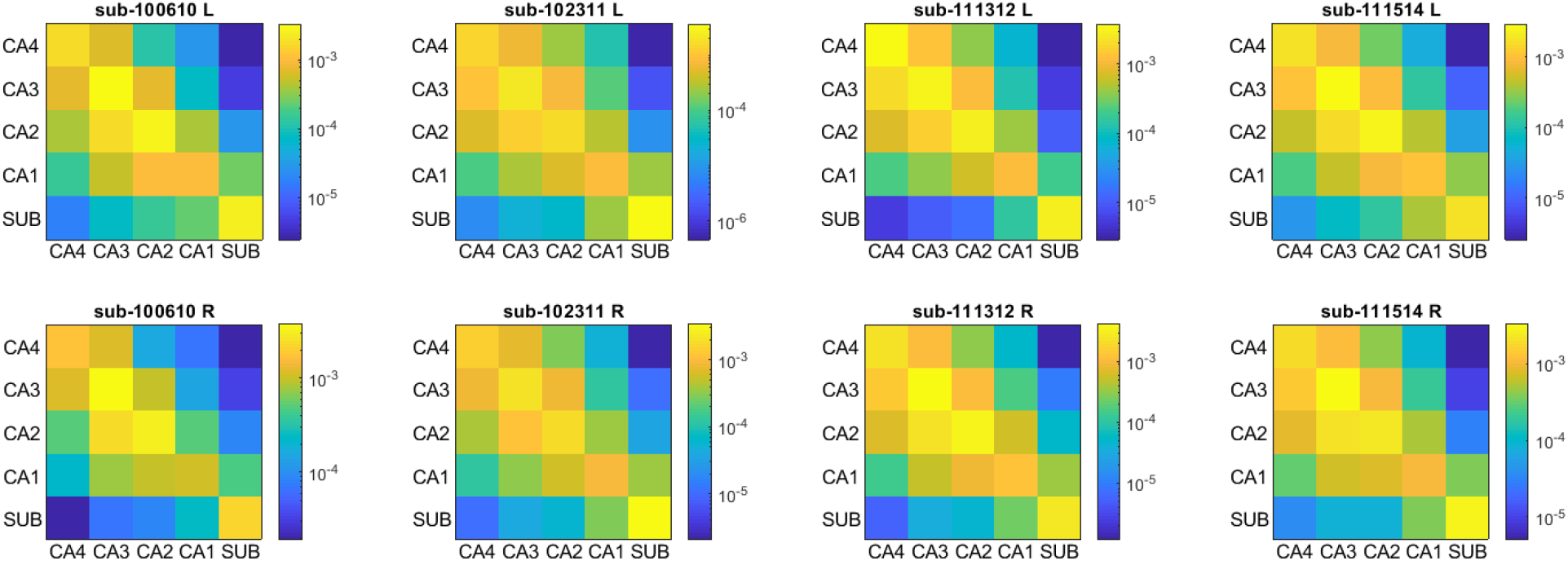
This figure shows the connectivity matrices *C_lm_* as calculated from `probtrackx` for the unfolded space at a resolution of 0.625 mm. The columns are the different participants and the top and bottom rows are the left and right hemispheres respectively.

**Figure 16:**
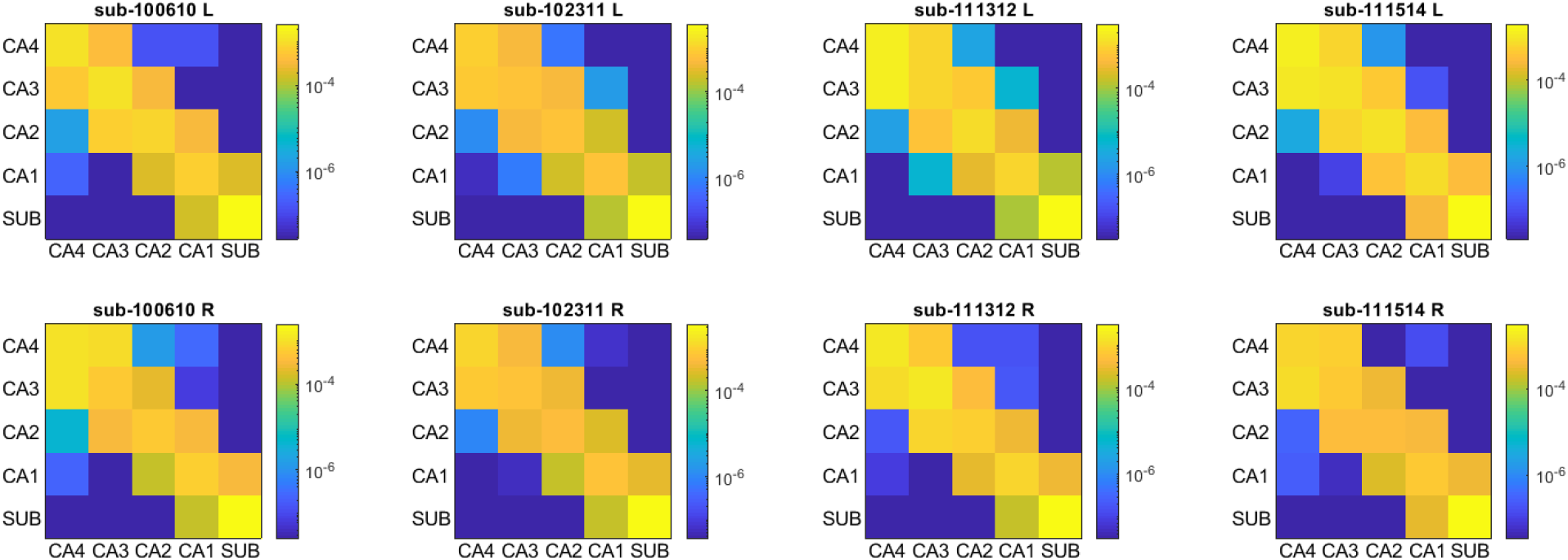
This figure shows the connectivity matrices 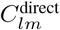 as calculated from `probtrackx` for the unfolded space at a resolution of 0.625 mm. The columns are the different participants and the top and bottom rows are the left and right hemispheres respectively.

