## Supplementary material for "Diffusion MRI of the Unfolded Hippocampus"

To make comparisons with trends observed with tracer injections in the nonhuman primate hippocampus, we filter the large connectivity matrix $\tilde{C}_{ij}$ to show the average connectivity between two regions in the unfolded hippocampus. This procedure is done by extracting rows in $\tilde{C}_{ij}$ that correspond to voxels in the first region and then we collapse these rows by taking an average. In this collapsed row vector we then extract columns that correspond to voxels in the second region and collapse to a scalar by taking an average. The figure below shows explicitly which regions we select when we perform our comparisons.

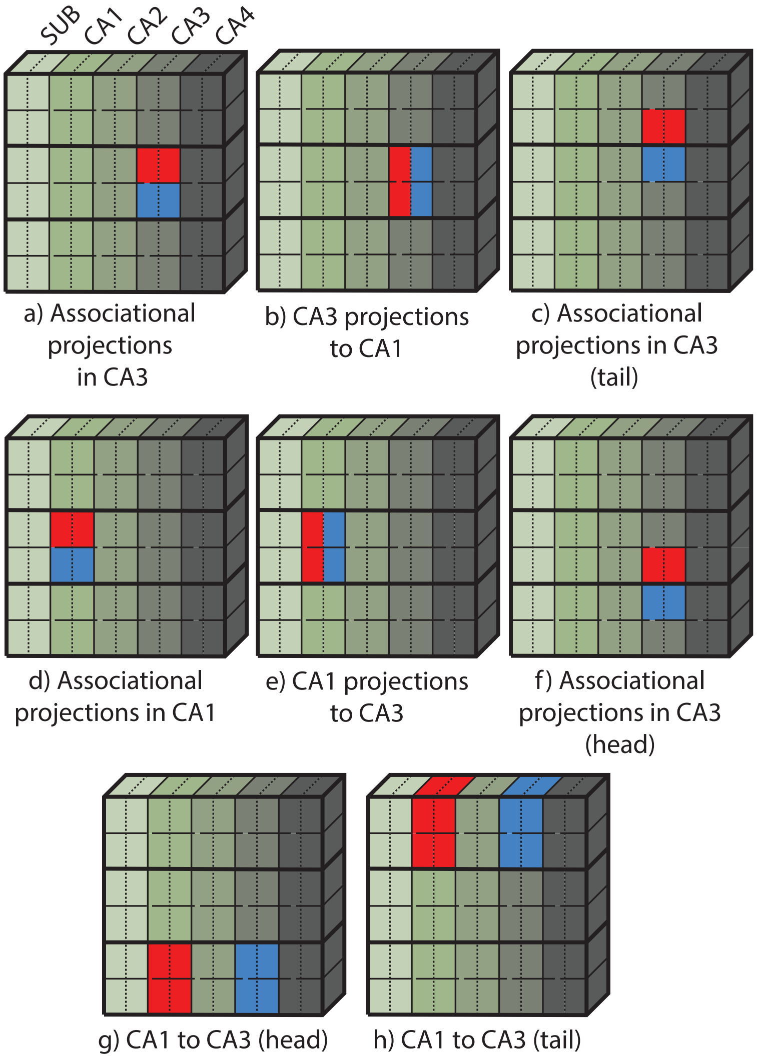


Figure 1: An illustration of the regions (red and blue) we select in the unfolded hippocampus to test the connectivity trends observed in the non human primate hippocampus through tracer injections. Note that we use b) when comparing with a), and e) when comparing with d).
